# Assessment of a split homing based gene drive for efficient knockout of multiple genes

**DOI:** 10.1101/706929

**Authors:** Nikolay P. Kandul, Junru Liu, Anna Buchman, Valentino M. Gantz, Ethan Bier, Omar S. Akbari

## Abstract

Homing based gene drives (HGD) possess the potential to spread linked cargo genes into natural populations and are poised to revolutionize population control of animals. Given that host-encoded genes have been identified that are important for pathogen transmission, targeting these genes using guide RNAs as cargo genes linked to drives may provide a robust method to prevent transmission. However, effectiveness of the inclusion of additional guide RNAs that target separate host encoded genes has not been thoroughly explored. To test this approach, here we generated a split-HGD in *Drosophila melanogaster* that encoded a drive linked effector consisting of a second gRNA engineered to target a separate host encoded gene, which we term a gRNA-mediated effector (GME). This design enabled us to assess homing and knockout efficiencies of two target genes simultaneously, and also explore the timing and tissue specificity of Cas9 expression on cleavage/homing rates. We demonstrate that inclusion of a GME can result in high efficiency of disruption of its target gene during super-Mendelian propagation of split-HGD. However, maternal deposition and embryonic expression of Cas9 resulted in the generation of drive resistant alleles which can accumulate and limit the spread of such a drive. Alternative design principles are discussed that could mitigate the accumulation of resistance alleles while incorporating a GME.

## Introduction

For standard Mendelian inheritance, any particular allele has a 50% chance in being transmitted to its offspring. While mechanisms of meiosis generally bias selection against violators of Mendel’s rules, there are many examples of naturally occurring selfish genetic elements (SGEs) that succeed in bypassing these rules. These SGEs enhance, or “drive” their transmission into subsequent generations, despite often times being harmful to the harboring individual (i.e. imposing a fitness load). These include, for example, transposable elements (TEs), meiotic drivers, B chromosomes, post segregation killers, heritable microbes, and homing endonuclease genes (Burt and Trivers, 2006; McLaughlin and Malik, 2017; Werren, 2011; Werren et al., 1988). Drawing inspiration from these natural systems, strategies for exploiting drive to alter the genetics of wild pest populations have been proposed (Burt and Trivers, 2006; Champer et al., 2016; Esvelt et al., 2014; McLaughlin and Malik, 2017; Werren, 2011; Werren et al., 1988), and some have even been experimentally tested in the laboratory, however none have been implemented in the field. For those tested in the laboratory, some examples include synthetic *Medea* elements (Akbari et al., 2014; A. Buchman et al., 2018; Chen et al., 2007), engineered underdominance systems (Akbari et al., 2013; A. B. Buchman et al., 2018), and those whose development was accelerated by the CRISPR revolution (Cong et al., 2013; Jinek et al., 2012; Mali et al., 2013) including toxin-antidote based systems (Oberhofer et al., 2019), and homing based gene drive systems (HGDs) (Champer et al., 2016; Esvelt et al., 2014; Gantz and Bier, 2016; Marshall and Akbari, 2018). HGDs are perhaps the furthest along in development, and in fact in addition to model systems these have already been tested in mosquitoes and even mammals (Champer et al., 2018, 2017; DiCarlo et al., 2015; Gantz et al., 2015; Gantz and Bier, 2015; Grunwald et al., 2019; Hammond et al., 2016, 2018; KaramiNejadRanjbar et al., 2018; Kyrou et al., 2018; Li et al., 2019; Windbichler et al., 2011; Yan and Finnigan, 2018). They function by encoding the Cas9 endonuclease and an independently expressed guide RNA (gRNA) responsible for mediating DNA/RNA base pairing and cleavage at a predetermined site (Champer et al., 2016; Esvelt et al., 2014; Gantz and Bier, 2016; Marshall and Akbari, 2018). When the HGD is positioned within its target site in a heterozygote, double stranded DNA breakage of the opposite chromosome can result in the drive allele being used as a template (i.e. donor chromosome) for DNA repair mediated by homologous recombination. This can result in copying, or “homing,” of the HGD into the broken chromosome (i.e. receiver chromosome), thereby converting heterozygotes to homozygotes in the germline, which can bias Mendelian inheritance ratios and result in an increase in HGD frequency in a population.

Given the recent progress toward developing HGDs in pest species such as mosquitoes (Gantz et al., 2015; Hammond et al., 2016, 2018; Kyrou et al., 2018; Li et al., 2019), there is significant enthusiasm regarding their potential use to control wild populations. For example, release of HGDs linked with effector genes inhibiting mosquito pathogen transmission (Buchman et al., 2019a, 2019b; Isaacs et al., 2011; Jupatanakul et al., 2017) may lead to replacement of disease-susceptible mosquitoes with disease-resistant counterparts resulting in reduced pathogen transmission (i.e. population modification drive). Alternatively, HGDs targeting genes affecting the fitness of female mosquitoes could also spread, resulting in gradual population declines and potentially even elimination (i.e. population suppression drive) (Kyrou et al., 2018; Windbichler et al., 2011, 2008). Given these features, both modification and suppression drives possess the potential to transform mosquito population control measures (Burt, 2003; Champer et al., 2016; Esvelt et al., 2014), and therefore have excited significant ongoing discussions involving their potential usage, regulation, safety, ethics and governance (Adelman et al., 2017; Akbari et al., 2015; National Academies of Sciences, Engineering, and Medicine et al., 2016; Oye et al., 2014). Despite these exciting developments however, the elephant in the room persists - can a gene drive actually work in the wild? There are a number of open questions looming as to the efficiency of a HGDs. For example, can a drive spread to fixation in the wild? Will it simply breakdown due to resistance? Will the linked anti-pathogen effector work efficiently given the expected diversity of parasites/virus genomes found in the wild? Can the pathogen evolve to become resistant to the anti-pathogen effector and perhaps even become more virulent? These are just a minority of legitimate concerns regarding the potential use of a gene drive that would need to be resolved prior to any release.

While many questions loom, there has been some effort to resolve these concerns safely in the lab. For example, with regard to the HGD breakdown due to resistance, multiple studies have explored design criteria attempting to suppress the effects of resistance alleles on drive propagation. For example, some studies have had some success using germline-restricted promoters to express Cas9 increasing rates of HDR, resulting in increased homing rates, as opposed to error-prone pathways such as non-homologous end joining (NHEJ) which results in the generation of resistant alleles (Champer et al., 2018; Hammond et al., 2018). Other studies have described (Champer et al., 2016; Esvelt et al., 2014; Marshall et al., 2017) and tested (Champer et al., 2018; J. Champer et al., 2019; S. E. Champer et al., 2019; Oberhofer et al., 2018) multiplexed gRNAs in drives resulting in moderate increases in drive efficacy. While others have had some success targeting highly conserved recessive fertility/viability genes whose homozygous mutants are inviable, or cannot reproduce, and therefore are expected to not affect the spread of HGDs (Hammond et al., 2016; KaramiNejadRanjbar et al., 2018; Kyrou et al., 2018; Oberhofer et al., 2018). However, despite these efforts, resistance alleles are still problematic, leaving open the question as to what is the best method to prevent their generation.

Here, to further explore this paramount issue of resistance to HGD we use *Drosophila melanogaster* as our model. We use a genetic safeguarded split-drive design as a safety feature and also encode a linked effector to the drive. This effector consisted of a second gRNA engineered to target a separate host encoded gene which we term a gRNA-mediated effector (GME) (Fig. S1A-B). Given that there are many host-encoded genes that are important for pathogen transmission (Cheng et al., 2016; Dong et al., 2018), one potential application of a HGD is to incorporate a cargo GME that targets a host encoded factor that is important for some aspect of pathogen transmission. If the GME is effective, then disruption of its target in the population should in principle occur as the drive spreads, thereby immunizing that population from pathogen transmission. Therefore, encoding a GME in a drive maybe useful feature going forward and worth further exploring. As a proof of concept to test the efficiency of a HGD linked GME, we designed both the drive and effector to target phenotypic genes which resulted in easily scorable recessive viable phenotypes. This novel drive architecture enabled us to test many germline Cas9 expressing promoters, while simultaneously measuring homing and cleavage efficiencies in both the germline and soma for both target genes over successive generations. While homing rates were modest, cleavage rates were high. For example, we determined that we can reproducibly achieve complete penetrance of somatic mosaic phenotypes for both target genes with up to 100% efficiency stemming from a combination of Cas9 maternal deposition and embryonic expression. However, despite the robust cleavage efficiencies and impressive efficacy of the HGD linked GME, drive resistance alleles were still generated which would hinder spread. Given these results, alternative design principles are proposed that could potentially mitigate these issues while also incorporating a drive linked GME (Fig. S2).

## Results

### Design of split-HGD encoding two gRNAs

To assess the feasibility and efficiency of utilizing a HGD to bias transmission while also expressing a GME, we designed a HGD that expressed two gRNAs (Lopez del Amo et al., 2019). The homing component of the split-HGD system, referred hereon as a Gene Drive element (GDe), encodes a gRNA targeting *white* (*gRNA*^*w*^, driver gRNA), a separate cargo GME targeting *yellow* (*gRNA*^*y*^, effector), a *3xP3-eGFP* dominant marker, all together flanked by 1kb homology arms from the *white* target locus to direct targeted HDR mediated integration (Fig. S1A). The *GDe* was integrated at the *white* locus (*w*^*GDe*^) in *D. melanogaster* via HDR (Fig. S1B). In the presence of Cas9, the *GDe* directs cleavage at both *white* and *yellow*, both X-linked loci, and is also capable of homing into the *white* locus (Fig. 1A). Importantly, in *D. melanogaster* homozygous loss-of-function (LOF) mutants of both *white* and *yellow* are viable and fertile with scorable recessive phenotypes in the eye and body, respectively, enabling cleavage events to be directly quantified over successive generations. Additionally, males have only one X chromosome, and are therefore hemizygous for *white* and *yellow*, restricting the quantification of homing to heterozygous females (y−,*w*^*GDe*^/*y*+,*w*+, Fig. 1A).

**Fig. 1.**
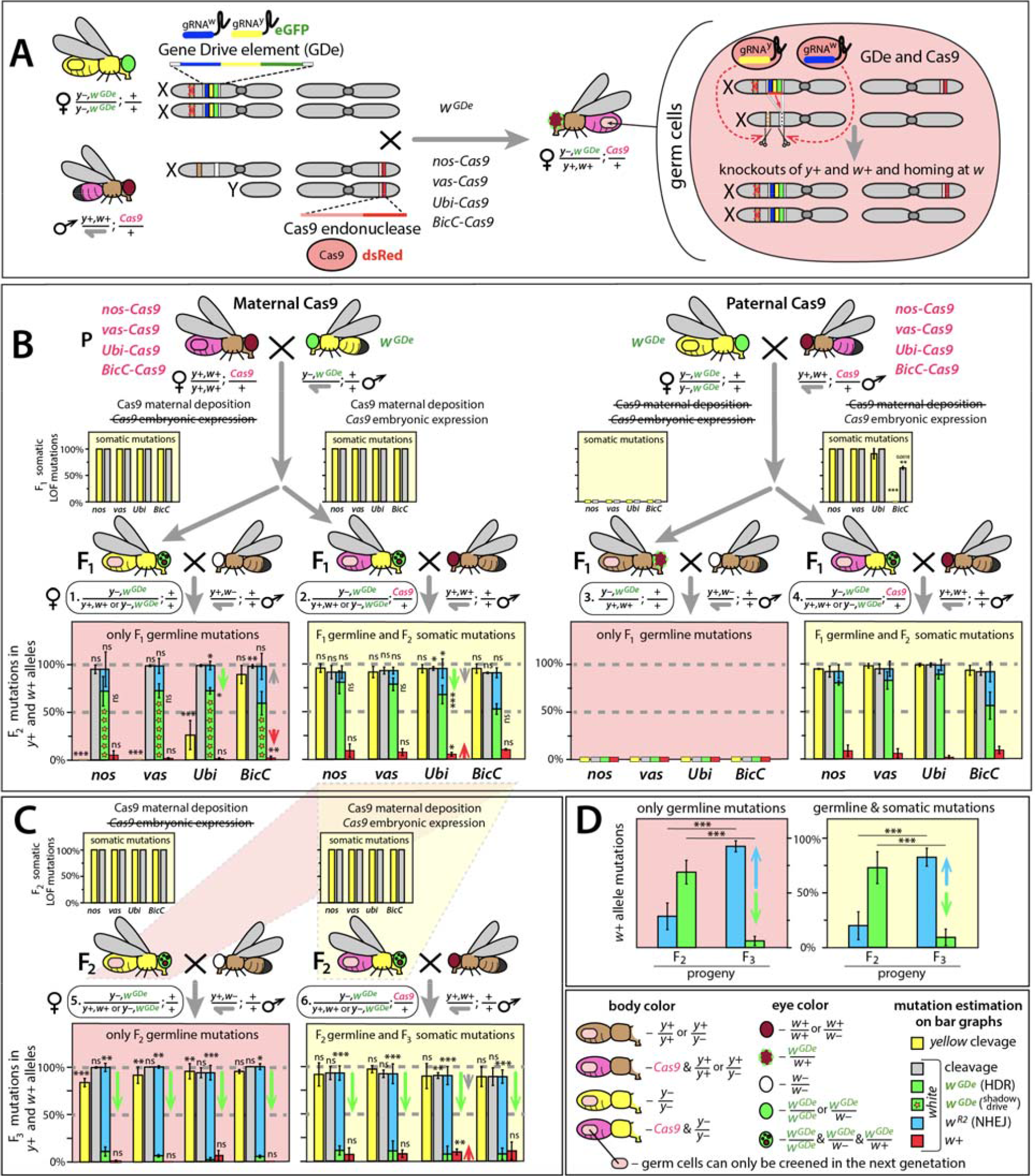
Split-HGD system forces super-Mendelian transmission and knocks out multiple genes. (**A**) Gene Drive element (GDe) with two gRNAs targeting *yellow* and *white* genes, and 3xP3-eGFP marker gene was interated via HDR at the *white* cut site on X sex chromosome (*w*^*GDe*^). *SpCas9* (Cas9) endonuclease expressed with one out of four promoters - *nanos* (*nos*), *vasa* (*vas*), *Bicaudal C* (*BicC*), and *Ubiquitin 63E* (*Ubi*) - and Opie2-dsRed marker gene (Fig. S1A) were integrated at the same site on the 3^rd^ chromosome. In heterozygous *y*−,*w*^*GDe*^/*y*+,*w*+; *Cas9*/+ females generated by a genetic cross between the *w*^*GDe*^ line and a *Cas9* line, *GDe* induces DSBs in *y*+ and *w*+ alleles resulting in cleavage of both genes and copying itself into the cut *w*+ allele via HDR (i.e. homing). The *w*^*GDe*^ allele cannot home in *Drosophila* males, because they have only one X chromosome, aka. hemizygous. (**B**) Each tested Cas9 promoter, including previously characterised germline-limited *nos*, *vas*, and *BicC* promoters, supported Cas9 expression in F_1_ somatic tissues and resulted in *white* and *yellow* loss-of-function (LOF) mutations in the F_1_. Furthermore, maternal deposition of Cas9 protein alone was sufficient to generate F_1_ somatic *white* and *yellow* LOF mutations as well as induce both homing (*w*^*GDe*^) and formation of resistance alleles (*w*^*R2*^) in *w*+ alleles of their germ cells (F_1_ ♀ #1). Paternal deposition of Cas9 protein did not induce mutations in somatic or germ cells (F_1_ ♀ #3). Notably, while 100% F_1_ parents had *yellow* LOF somatic mutations with each tested Cas9 line, only *Ubi* and *BicC* promoters deposited Cas9 protein sufficient to induce *yellow* LOF mutations in some germ cells (F_1_ ♀ #1). Embryonic expression of *Cas9* gene under *nos*, *vas*, and *Ubi* promoters induced *white* and *yellow* LOF mutations in 100% trans-heterozygous females, while embryonic expression of *BicC-Cas9* caused only *white* LOF mutation in 64.3% ± 2.6% of trans-heterozygous females (F_1_ ♀ #4). Rates of homing and resistance alleles were not significantly different among two types of trans-heterozygous (F_1_ ♀ #2 and #4) and heterozygous (♀ #1) females with maternally deposited Cas9. Only maternal deposition of Cas9 under *Ubi* promoter negatively affected homing rates (green arrows) in germ cells. (**C**) Resistance alleles are expected to be immune to the further cleavage by the same Cas9/gRNA system and if their carrier are fertile can propagate at the expense of homing. To explore this idea, F_2_ ♀ #5 and F_2_ ♀ #6 collected among progeny of F_1_ ♀ #2 were genetically crossed with *w*− and *w*+ males, respectively, and their F_3_ progeny were scored. While the cleavage rate in F_2_ germ cells decreased only in F_2_ ♀ #6 with *Ubi-Cas9* (red arrow) likely due to the rise of functional w^*R1*^ alleles, the homing frequency fell significantly for each tested split-HGD system with and without *Cas9* gene (green arrows). The fall of homing rate was accompanied by the accumulation of the *w*^*R2*^ alleles. (**D**) Accumulation of *w*^*R2*^ alleles resistant to cleavage by Cas9/gRNA^w^ suppressed homing of GDe. Frequencies of homing and resistance alleles were averaged for all tested promoters and presented separately for progeny of heterozygous and trans-heterozygous females, F_2_ ♀ #5 and F_2_ ♀ #6, respectively. Resistance allele frequency increase from 28.5% or 19.9% to 92.6% or 82.6% between F_2_ and F_3_ (blue arrows) and caused the dramatic decline in homing from 69.0% or 73.0% to 6.1% or 9.2% (green arrows). Notably, scoring of *w*^*R2*^ alleles in *w*− recessive background resulted in the higher estimation of *white* LOF mutations alleles, since *w*^*R2*^ alleles were complemented by *w*+ alleles inherited from wild type males. Bar plots show the average ± SD over at least three biological replicate crosses. Statistical significance was estimated using a *t* test with equal variance. (*P* ≥ 0.05^ns^, *P* < 0.05*, *P* < 0.01**, and *P* < 0.001***).

### High penetrance of F_1_ somatic mutations generated by Cas9 through both maternal deposition and embryonic expression

To explore the effects of tissue specificity and timing of Cas9 expression on cleavage and homing in the germline, we used four separate promoters with distinct expression profiles to express Cas9-T2A-GFP: *nanos (nos)* (Van Doren et al., 1998) and *vasa (vas)* promoters known for early germline-limited expression (Hay et al., 1988; Sano et al., 2002; Van Doren et al., 1998); *Bicaudal C* (*BicC)* promoter supporting later germline-limited expression (Saffman et al., 1998); and *Ubiquitin 63E* (*Ubi*) promoter with strong expression in both somatic and germ cells (Akbari et al., 2009; Preston et al., 2006). To control for position effect variegation (PEV), each Cas9 construct (Fig. S1A) was integrated site specifically on the 3^rd^ chromosome using φC31-mediated integration (Groth, 2004). To confirm germline expression, we imaged the expression of a self-cleaving T2A-eGFP tag attached to the coding sequence of Cas9, and each promoter robustly expressed GFP in the ovaries (Lowest -Nanos-Cas9 < Vasa-Cas9 <Ubi-Cas9 <BicC-Cas9 Highest). (Fig. S3).

To quantify cleavage efficiencies, we performed bi-directional crosses between hemizygous or homozygous GDe lines mated to heterozygous Cas9 lines (Fig. 1B). From these crosses we determined that maternally deposited Cas9 protein is sufficient to induce both *yellow* and *white* somatic LOF mutations in F_1_ females heterozygous for the *GDe* both in presence (♀# 2; *y*−,*w*^*GDe*^/*y*+,*w*+; Cas9/+; Fig. 1B) and in the absence (♀ # 1; *y*−,*w*^*GDe*^/*y*+,*w*+; Fig. 1B) of *Cas9* gene inheritance. To determine whether embryonic expression of *Cas9* can also induce somatic mutations, we scored *white* and *yellow* LOF somatic mutations in F_1_ transheterozygous females inheriting *Cas9* exclusively from their fathers (i.e. paternal *Cas9*). Unexpectedly, F_1_ transheterozygous female progeny inheriting Cas9 as a gene (♀#4; y−,*w*^*GDe*^/*y*+,*w*+; Cas9/+; Fig. 1B) from their fathers had mutations in both *white* and *yellow* with varying frequencies depending on which promoter drove Cas9 expression. For example, *nos-Cas9* and *vas-Cas9* - induced 100% *white* and *yellow* LOF somatic mutations in F_1_ transheterozygous females, while *Ubi-Cas9* resulted in 100% of *white* and 91.3% ± 9.7% of *yellow* LOF somatic mutations, and *BicC-Cas9* resulted in only *white* LOF mutations in 64.3% ± 2.6% of the F_1_ y−,*w*^*GDe*^/*y*+,*w*+; *BicC-Cas9*/+ progeny (Fig. 1B). Interestingly however, 100% of F_1_ heterozygous female progeny from the same fathers that did not inherit Cas9 as a gene (♀#3; y−,*w*^*GDe*^/*y*+,w+; +/+; Fig. 1B) had wild type (*wt)* phenotypes, for both *white* (red eyes) and *yellow* (brown body), presumably resulting from lack of sufficient Cas9 protein deposited paternally to induce mutations in the zygote (Table S1). Taken together these data strongly indicate that the Cas9 promoters tested here are highly active both maternally and embryonically and can promote very high cleavage efficiency.

To determine whether *yellow* and *white* alleles were mutated by maternally deposited Cas9 in germ cells of F_1_ *y*−,*w*^*GDe*^/*y*+,*w*+ females, we mated these females to *y*+,*w*− males and scored recessive *yellow* phenotypes in resulting F_2_ male progeny (*y*−) and recessive *white* phenotypes in resulting F_2_ male and female progeny (*w*−/*w*−). Interestingly, we found that maternally deposited Cas9 protein expressed under *nos* and *vas* promoters did not induce *yellow* LOF mutations in germ cells of F_1_ females, while expression from *Ubi* and *BicC* promoters resulted in 26% ± 15% and 89.4% ± 9.4% of *yellow* alleles being mutated in germ cells of F_1_ females, respectively, perhaps due to a stronger maternal deposition of Cas9 protein by these promoters (Fig. S3) combined with possible prefentail gRNA loading by Cas9 (Fig. 1B). Despite the lack of LOF germline mutations in *yellow* by *nos* and *vas*, every tested Cas9 line provided a sufficient amount of maternally deposited Cas9 protein to knockout the *white* allele in 94.9% ± 4.5% to 98.8% ± 1.1% of F_1_ germ cells (measured in F_2_ progeny). To molecularly confirm whether the *w*+ alleles (1.2% - 5.1%) were cut by Cas9 and perhaps repaired into cleavage resistant alleles, we Sanger sequenced PCR amplicons of the *white* target locus from individual male flies. Each tested F_2_ male with red eyes (w+) indeed had a *wt w*+ allele, however we did not find any *white* in-frame functional resistant alleles in F_1_ germ cells suggesting that these alleles were likely remained uncut in the germline.

### Maternally deposited Cas9 is sufficient to induce homing of GDe in germ cells

The Cas9/gRNA^w^-induced DSBs at *white* locus can be repaired either by HDR resulting in homing of the *GDe* (*w*^*GDe*^/*w*^*GDe*^) or NHEJ incorporating *indel* mutations that can render the target locus unrecognizable by the Cas9/gRNA^w^ machinery, and when these mutations occur in germ cells they are referred to as resistance alleles (*w*^*R*^): here LOF and in-frame functional resistance alleles are referred as *w*^*R2*^ and *w*^*R1*^, respectively (Fig. S1C). To directly estimate the frequency of *w*^*GDe*^ homing and *w*^*R*^ generation in the absence of additional somatic mutations resulting from embryonic expression of *Cas9*, we analyzed *white* phenotypes in the F_2_ progeny of the F_1_ *w*^*GDe*^/*w*+ females with maternally deposited Cas9 in a *w*− recessive mutant background (Fig. 1B). Every tested Cas9 promoter provided a sufficient amount of maternally deposited Cas9 in the F_1_ germ cells to enable the conversion of 59% - 72% of *w*+ alleles into *w*^*GDe*^ (i.e. homing of *GDe)* in *y*−,*w*^*GDe*^/*y*+,*w*+ females. This conversion which occurs in the presence of Cas9 protein, but absence of inheritance of the *Cas9* gene, was previously noted and termed “shadow drive” (Guichard et al., 2019). The remaining DSBs at *w*+ alleles were repaired by NHEJ and generated around 38% - 23% *w*^*R2*^ alleles (Fig. 1B). To explore molecular changes at *white* locus, we PCR amplified and Sanger sequenced *w*^*R2*^ alleles from individual F_2_ male progeny and identified *indels* localized at the *white* cut site in each sequenced male (Fig. S4A). The maternally deposited Cas9 by *BicC* promoter resulted in the lowest homing and the highest resistance allele rates (59.3% ± 12.3% and 38.7% ± 13.7%, respectively), though no significant difference was identified between each pairwise comparison. Nevertheless, each tested promoter supplied Cas9 protein to the progeny that enabled shadow drive, and thus resulted in super-Mendelian propagation of *w*^*GDe*^ in their grandchildren.

### Maternal deposition of Cas9 protein reduces the homing efficiency

Maternally deposited Cas9 can induce *white* cleavage and repair mediated by NHEJ as opposed to HDR in mitotically dividing germ cells which can result in a bias toward generating resistance alleles (*w*^*R2*^ and *w*^*R1*^) at the expense of homing *w*^*GDe*^. To explore this effect, we compared homing rates between F_1_ trans-heterozygous females that inherited Cas9 either maternally (♀#2; *y*−,*w*^*GDe*^/*y*+,*w*+; Cas9/+) or paternally (♀#4; *y*−,*w*^*GDe*^/*y*+,*w*+; Cas9/+; Fig 1B). For *nos-Cas9, vas-cas9*, and *BicC-Cas9*, maternal deposition of Cas9 did not result in a significant bias in homing efficiencies. However, for *Ubi-Cas9* homing rates were significantly lower (67%) in the trans-heterozygous females that inherited *Cas9* maternally (♀#2; *w*^*GDe*^/*w*+; *Ubi-Cas9*/+) as compared to 88% for trans-heterozygous females inheriting *Ubi-Cas9* paternally (♀#4; *w*^*GDe*^/*w*+; *Ubi-Cas9*/+; Fig. 1B). In addition to the lower homing rates for *Ubi-Cas9*, the rate of *w*^*R2*^ alleles was significantly higher with maternally deposited *Ubi-Cas9* as compared to paternally deposited *Ubi-Cas9*: 9.9% ± 5.7% vs 27.3% ±10.0%, *P* > 0.025, vs 26.5% ± 4.4%, *P* > 0.029, respectively (Fig. 1B). Taken together, these results suggests that strong maternal deposition of Cas9 protein into developing oocytes can result in *white* cleavage in mitotic cells, prior to developmental stages where efficient HDR repair occurs, therefore leading to a higher *w*^*R*^ frequency.

### Resistance alleles accumulate between F_2_ and F_3_ generations

Resistance alleles generated in germ cells are immune to subsequent cleavage by the Cas9/gRNA^w^ complex. *Drosophila white* and *yellow* LOF homozygotes are viable and fertile, as a result, the frequency of resistance alleles can potentially increase from generation to generation. To explore this possibility, we crossed F_2_ trans-heterozygous females (♀ #6, *w*^*GDe*^/*w*+; *Ubi-Cas9*/+; Fig. 1C) to *wt* (*y*+,w+) males, and scored their F_3_ progeny for *yellow* and *white* phenotypes, as well as for inheritance of the *GDe*. Indeed, the frequency of *white* LOF mutations (*w*^*R2*^) increased significantly between F_2_ and F_3_ progenies for each Cas9 promoter: 11.2% ± 6.2% vs 81.7% ± 7.5% for *nos-Cas9*; 13.2% ± 5.6% vs 82.4% ± 10.4% for *vas-Cas9*; 18.6% ± 12.0% vs 84.6% ± 9.5% for *Ubi-Cas9*; and 36.7% ± 7.5% vs 81.6% ± 7.1% for BicC-Cas9, *P* > 0.0001, respectively. This rise of *w*^*R2*^ frequency negatively affected the homing rate, as it plummeted between F_2_ and F_3_ generations: from 80.0% ± 7.7% to 11.3% ± 4.8% for *nos-Cas9*; from 80.2% ± 7.4% to 10.8% ± 10.4% for *vas-Cas9*; from 78.0% ± 13.2% to 7.4% ± 8.4% for *Ubi-Cas9*; and from 53.9% ± 9.8% to 7.6% ± 6.2% for *BicC-Cas9* (♀ #6, *w*^*GDe*^/*w*+; *Ubi-Cas9*/+; Fig. 1C). To avoid any ambiguity caused by somatic expression of Cas9, the same analysis was repeated with the F_2_ heterozygous females carrying maternally deposited Cas9 protein but lacking the *Cas9* gene resulting in similar conclusions (♀ #5, *y*−,*w*^*GDe*^/*y*+,*w*+; Fig. 1C). To assess the accumulation of resistance alleles, we compared mean frequencies of homing and resistance alleles between F_2_ and F_3_ generations. The frequency of resistance alleles rose from 28.5% ± 12.2% to 92.6% ± 5.0% in heterozygous females or from 19.9% ± 12.8% to 82.6%±8.2% in trans-heterozygous females, and resulted in the decrease of homing rate from 69.0% ± 10.8% to 6.1% ± 4.2% or from 73.0%±14.6% to 9.2% ± 7.5%, respectively (*P* > 0.0001, Fig. 1D). As expected, the frequency of LOF resistance alleles at *white* locus (*w*^*R1*^) also increased from F_2_ to F_3_ progenies and restricted further homing of the *GDe*. The frequency of in-frame functional *white* and *yellow* mutations (*w*^*R1*^ and *y*^*R1*^) could also increase in the F_3_ progeny, but unfortunately it could not be directly estimated. The *white* cleavage frequency significantly decreased in the F_3_ progeny of F_2_ *y*−,*w*^*GDe*^/*y*+,*w*+; *Ubi-Cas9*/+ females, and could be explained by the increase of *w*^*R1*^ allele rate that were indistinguishable from *w*+ alleles phenotypically: from 3.4% ± 2.6% in F_2_ to 9.7% ± 3.1% in F_3_, *P* > 0.004 (Fig. 1B-C). To further explore this possibility, we Sanger sequenced F_3_ *wt* males with red eyes and brown bodies, and identified in-frame *indels* and substitutions in the majority of tested males for each Cas9 promoter (*w*^*R1*^ and *y*^*R1*^ alleles, Fig. S4). Therefore, many germ cells of F_2_ trans-heterozygous and heterozygous with maternally deposited Cas9 females has *indel* mutations at *white* and *yellow* loci (*y*−,*w*^*GDe*^/*y*^*R*^,*w*^*R*^) that were indeed resistant to further cleavage by Cas9/gRNA^w^ and Cas9/gRNA^y^, respectively.

## Discussion

Homing based gene drives require efficient cleavage and copying in the germline in order to bias their transmission and are therefore sensitive to both existing and induced target sequence variation. In fact, the NHEJ-mediated generation of resistance alleles in germ cells was previously identified as the major force opposing the spread of HGD into populations (Champer et al., 2017; Gantz et al., 2015; Hammond et al., 2017; Oberhofer et al., 2018). Here, in a split-drive design we further explored the effect of timing and expression of Cas9 on both homing and cleavage efficiencies. Additionally, we linked a GME to the drive to measure the efficacy of this approach. This drive architecture enabled us to draw several conclusions including; i) expression of a drive mediating gRNA in addition to a linked GME can result in 100% penetrance of both scorable LOF phenotypes; ii) each tested Cas9 expression promoter (*nos*, *vas*, *BicC*, Ubi) also results in significant embryonic expression; iii) Cas9 maternal protein deposition or embryonic expression are both sufficient for homing; iv) maternal deposition of Cas9 protein is not required for homing in females; v) paternal Cas9 protein deposition in the sperm is insufficient for either homing or cleavage; vi) resistant alleles accumulate over subsequent generations which are predicted to impair the spread of the drive. Below we discuss these conclusions further and also propose novel drive architectures to potentially overcome these issues.

### Somatic expression of Cas9 results in high mutagenesis rates

The maternal deposition and embryonic expression of *Cas9* in the presence of a *gRNA* transgene were previously shown to induce LOF mutations in F_1_ progeny from a cross using *nos*- or *vas*-driven *Cas9* and U6-*gRNA* lines (Kandul et al., 2019; Lin and Potter, 2016; Oberhofer et al., 2018; Port et al., 2014); however, the somatic nature of F_1_ LOF mutations was not fully explored. This is in part due to the fact that when Cas9 and gRNA are linked together in the single-locus HGD, somatic and germline LOF mutations are not easily distinguishable from heritable mutations occuring in prior generations which can result in overestimation of homing rates. Therefore, unlinking these components enables a better method for carefully distinguishing these events. Here, using a split-drive design, we were able to carefully assess the effects of timing, expression and inheritance of Cas9 on both homing and cleavage efficiencies. As reported previously, we found that maternal Cas9 protein deposition was sufficient to induce shadow drive (Guichard et al., 2019) in addition to high rates of F_1_ somatic LOF mutations (Kandul et al., 2019; Lin and Potter, 2016; Oberhofer et al., 2018; Port et al., 2014). However, unexpectedly we found that embryonic expression of Cas9 was also sufficient to induce cleavage and homing. In fact, both *nos* and *vas* promoters, which were previously characterized by early germline-limited expression (Sano et al., 2002; Van Doren et al., 1998), do support significant embryonic expression of Cas9. For example, F_1_ progeny with either maternally deposited Cas9 protein, or embryonically expressed *Cas9* gene inherited from their fathers, both had moderate rates of HDR in addition to *white* and *yellow* LOF somatic mutations with up to 100% efficiency (Fig. 1B); consistent observations were generated in a recent work using a trans-complementing Gene Drive (tGD) system (Lopez del Amo et al., 2019). These data conclusively demonstrate that Cas9 is embryonically expressed and that embryonic expression is sufficient to generate high rates of cleavage and moderate rates of HDR. Given this high penetrance of cleavage, this data suggest that effectors that mechanically rely on cleavage (i.e. GME’s) could indeed be quite effective if linked to an efficacious drive.

### Resistant alleles accumulate over subsequent generations

Consistent with previous studies, we also found that maternal deposition of Cas9 protein into the embryo can result in both cleavage and homing in the germ cells (Guichard et al., 2019). In addition to this observation, we also found that paternal Cas9 protein deposition was not sufficient to induce either cleavage or homing, presumably due to the low quantities of Cas9 carried by the sperm into the egg. Moreover, we determined that maternal deposition is not a requirement for homing, and in fact females that inherit the *Cas9* gene paternally can indeed express Cas9 embryonically resulting in both cleavage and homing. Notwithstanding, regardless of whether *Cas9* was maternally or paternally inherited, there was no significant difference between the rates of appearance of drive resistance alleles which were generated at moderate frequencies in the F_2_. Interestingly, the frequency of resistance alleles (*w*^*R*^) increased dramatically between F_2_ and F_3_ generations and correlated with decreases in homing (Fig. 1D). Taken together, these results suggest that homing occurs post embryonically and requires sufficient Cas9 to be present in the egg via either maternal deposition, or embryonic expression, both of which lead to the generation and accumulation of resistant alleles predicted to impede the spread of the drive.

### Novel strategies for disarming resistant alleles in germ cells

The accumulation of drive resistant alleles reported here was in part due to the fact that *white* is recessive viable, enabling targeted drive resistant alleles to accumulate. Given this accumulation, perhaps targeting non-essential genes using HGD may not be ideal. To avoid this issue, targeting essential genes would be a more appropriate design to ensure gene drive stability and spread. By targeting essential genes, it is possible that non-drive alleles could be actively selected against using a phenomenon previously termed lethal mosaicism (Guichard et al., 2019; Kandul et al., 2019) or by natural selection due to increased fitness costs. Lethal mosaicism results in dominant biallelic knockouts of target genes throughout development which could eliminate cleavage resistant alleles as they would be non-viable. We envision two novel drive design architectures that incorporate a GME and rely on lethal mosaicism to limit the generation of resistant alleles. First, haplo-sufficient genes essential for insect viability or fertility can be targeted by HGD designed to express a recoded version of the disrupted gene that is resistant to gRNA-mediated cleavage in addition to a linked GME (HGD+R+GME). This ensures that only the progeny that inherit the HGD+R+GME survive, while all progeny that inherit a cleaved allele perish due to non-rescued lethal mosaicism (Fig. S2). Second, a **<u>c</u>**leavage-only **<u>g</u>**ene **<u>d</u>**rive with rescue could be designed that incorporates a GME (CGD+R+GME) which mechanistically relies exclusively on cleavage for biased inheritance and selection against drive resistant alleles (Fig. S2). Both of these strategies would likely be effective to limit the accumulation of drive resistance alleles. However, in-frame functional mutations (*R1* type) that confer resistance against the Cas9/gRNA and do not cause fitness costs carriers may still be generated which could limit the spread of a drive. To summarize, our results demonstrate that inserting a GME into a HGD efficient knockouts of multiple genes can be achieved while simultaneously biasing *GDe* transmission rates into subsequent generations. However, resistant alleles were generated, and accumulated, which would limit the efficacy and spread of this system. To overcome these limitations, novel drive architectures are proposed and remain to be tested in future studies.

## Materials and methods

### Design and assembly of constructs

The genetic assembly of the Gene Drive element with gRNAs and 3xP3-eGFP (*GDe*) and employment of the *w*^*GDe*^/*w*^*GDe*^ which was previously used in for a different purpose by (Lopez del Amo et al., 2019) to generate a split trans-complementing Gene Drive system. The assembly of *BicC-Cas9* construct followed the same steps previously described for the other three Cas9 lines: *nos-Cas9*, *vas-Cas9*, and *Ubi-Cas9* (Kandul et al., 2019). The 2831 bases upstream of BicC-RA’s start codon (*Bicaudal C*, CG4824) was PCR amplified with CGACGGTCACGGCGGGCATGTCGACGCGGCCGCATAATTATATAATAATAAACTGC ATGC (BicC-F) and TCCGTCGTGGTCCTTATAGTCCATGTTTAAACTGTGGAATTCGGATGATGATGATGATC (BicC-R) from *Drosophila melanogaster* genome, and enzymatically assembled (Gibson et al., 2009) into *Ubi-Cas9* plasmid (addgene #112686) (Kandul et al., 2019) digested with NotI and XhoI.

### Fly genetics and imaging

Flies were maintained under standard conditions at 25□°C. Embryo injections were carried at Rainbow Transgenic Flies, Inc. (http://www.rainbowgene.com). The *BicC-Cas9* construct was inserted at the PBac{y+-attP-3B}KV00033 on the 3^rd^ chromosome (Bloomington #9750) with φC31-mediated integration (Groth, 2004). Transgenic flies were balanced with Df(3L)R/TM6C,cu^1^,Sb^1^,Tb^1^ (Bloomington #57) and CxD,ry^BM^/TM3, Sb^1^,Ser^1^ (Bloomington #1704) in the *w*+ genetic background.

To assess the cleavage rates and homing efficiencies of the split-HGD system, we genetically crossed the *GDe* line to four different *Cas9* lines in both directions. Two types of F_1_ trans-heterozygous *y−,w^GDe^(eGFP)/y+,w+*; *Cas9(RFP)*/+ females carrying either maternal or paternal *Cas9* (F_1_ ♀ #2 or ♀ #4, respectively) and the F_1_ heterozygous *y*−,*w*^*GDe*^(eGFP)/*y*+,*w*+ females with maternally deposited Cas9 were generated (F_1_ ♀ #1, Fig. 1B). Their *yellow* and *white* LOF mutations and transgene markers were scored. To explore whether *yellow* and *white* loci were also mutated in the germ cells of the F_1_ trans-heterozygous and heterozygous females, we genetically crossed them to *w*+,*y*+ and *w*-,*y*+ males, respectively, and examined their F_2_ progeny. LOF *yellow* mutations were scored only in male progeny that inherited their single X chromosome from mothers. To explore the behaviour of resistance alleles over multiple generations, the F_2_ trans-heterozygous and heterozygous virgin female (♀ #6 or ♀ #5, respectively) progeny of F_1_ ♀ #2 were also collected, and genetic crosses and phenotype scoring were repeated for an additional generation, F_3_. The above crossing schemes are depicted in Fig. 1B. To generate means and standard deviations for statistical comparisons, each genetic cross was set up in triplicate using 10♂ and 10♀ flies for each replicate cross. Cleavage and homing frequencies are presented as percentages of *y*+ and *w*+ alleles in heterozygous females, aka. they normalized to 50% (Table S1)

Flies were examined, scored, and imaged on the Leica M165FC fluorescent stereo microscope equipped with the Leica DMC2900 camera. To analyze Cas9 expression in ovaries of four homozygous *Cas9* lines, their ovaries were dissected in PBS buffer, examined, and imaged utilizing the same settings. The eGFP fluorescence was used as a proxy of Cas9 expression, since it was tagged to *Cas9* transgene as via a *T2A* sequence (Fig. S3).

### Genotyping loci targeted with gRNAs

To explore the molecular changes that caused LOF and in-frame functional mutations in *yellow* and *white* loci, we PCR amplified the genomic regions containing target sites for *gRNA*^*w*^ and *gRNA*^*y*^: GGCGATACTTGGATGCCCTGCGG and GGTTTTGGACACTGGAACCGTGG, respectively. Single-fly genomic DNA preps were prepared by homogenizing a fly in 30μl of a freshly prepared squishing buffer (10mM Tris-Cl pH 8.0, 1mM EDTA, 25mM NaCL, 200 μg/mL Proteinase K), incubating at 37°C for 35 minutes, and heating at 95°C for 2 minutes. 2 μl of genomic DNA was used as template in a 40 μL PCR reaction with LongAmp® Taq DNA Polymerase (NEB). The 415bp PCR fragment of *white* target was amplified with CGTTAGGGAGCCGATAAAGAGGTCATCC (w.sF) and AAGAACGGTGAGTTTCTATTCGCAGTCGG (w.sR); and CACTCTGACCTATATAAACATGGACCGCAGTTTG (y.sF) and CCAATTCATCGGCAAAATAGGCATATGCAT (y.sR) primers were used to amplify the 375bp PCR fragment of *yellow*. PCR aplicons were purified using QIAquick PCR purification kit (QIAGEN), and sequenced in both directions with Sanger method at Source BioScience. To characterize molecular changes at the targeted sites, sequence AB1 files were aligned against the corresponding reference sequences in SnapGene® 4.

### Statistical analysis

Statistical analysis was performed in JMP 8.0.2 by SAS Institute Inc. At least three biological replicates were used to generate statistical means for comparisons. To estimate the effect of Cas9 maternal deposition on homing efficiency, rates of cleavage, homing, and resistance alleles in F_1_♀ #4 with paternal Cas9 were compared to the corresponding values in F_1_ ♀ #1 and ♀ #2 with maternally deposited Cas9 protein (Fig. 1B). To assess the significance of resistance allele accumulation and homing rate decline between F_2_ and F_3_ generations, rates of cleavage, homing, and resistance alleles in F_2_ ♀ #5 and F_2_ ♀ #6 (Fig. 1C) were compared to the corresponding values in F_1_ ♀ #1 and F_1_ ♀ #2, respectively (Fig. 1B). *P* values were calculated for a two-sample Student’s *t*-test with equal variance.

### Gene Drive safety measures

All crosses using gene drives genetics were performed in accordance to an Institutional Biosafety Committee-approved protocol from UCSD in which full gene-drive experiments are performed in a high-security ACL2 barrier facility and split drive experiments are performed in an ACL1 insectary in plastic vials that are autoclaved prior to being discarded in accord with currently suggested guidelines for laboratory confinement of gene-drive systems (Akbari et al., 2015; National Academies of Sciences, Engineering, and Medicine et al., 2016).

### Ethical conduct of research

We have complied with all relevant ethical regulations for animal testing and research and conformed to the UCSD institutionally approved biological use authorization protocol (BUA #R2401).

## Acknowledgements

This work was supported in part by the University of California, San Diego, Department of Biological Sciences, by funding from the Defense Advanced Research Project Agency (DARPA) under a “Safe Genes” Program Grant (HR0011-17-2-0047) awarded to O.S.A. and by the Office of the Director of the National Institutes of Health under award number DP5OD023098 awarded to V.M.G.

## Author Contributions

O.S.A, and N.P.K. conceptualized the study. A.B. and V.M.G. designed and assembled constructs for this project. N.P.K. and J.L. performed molecular and genetic experiments. All authors contributed to the writing, analyzed the data, and approved the final manuscript.

---

**Supplementary Fig. S1.**
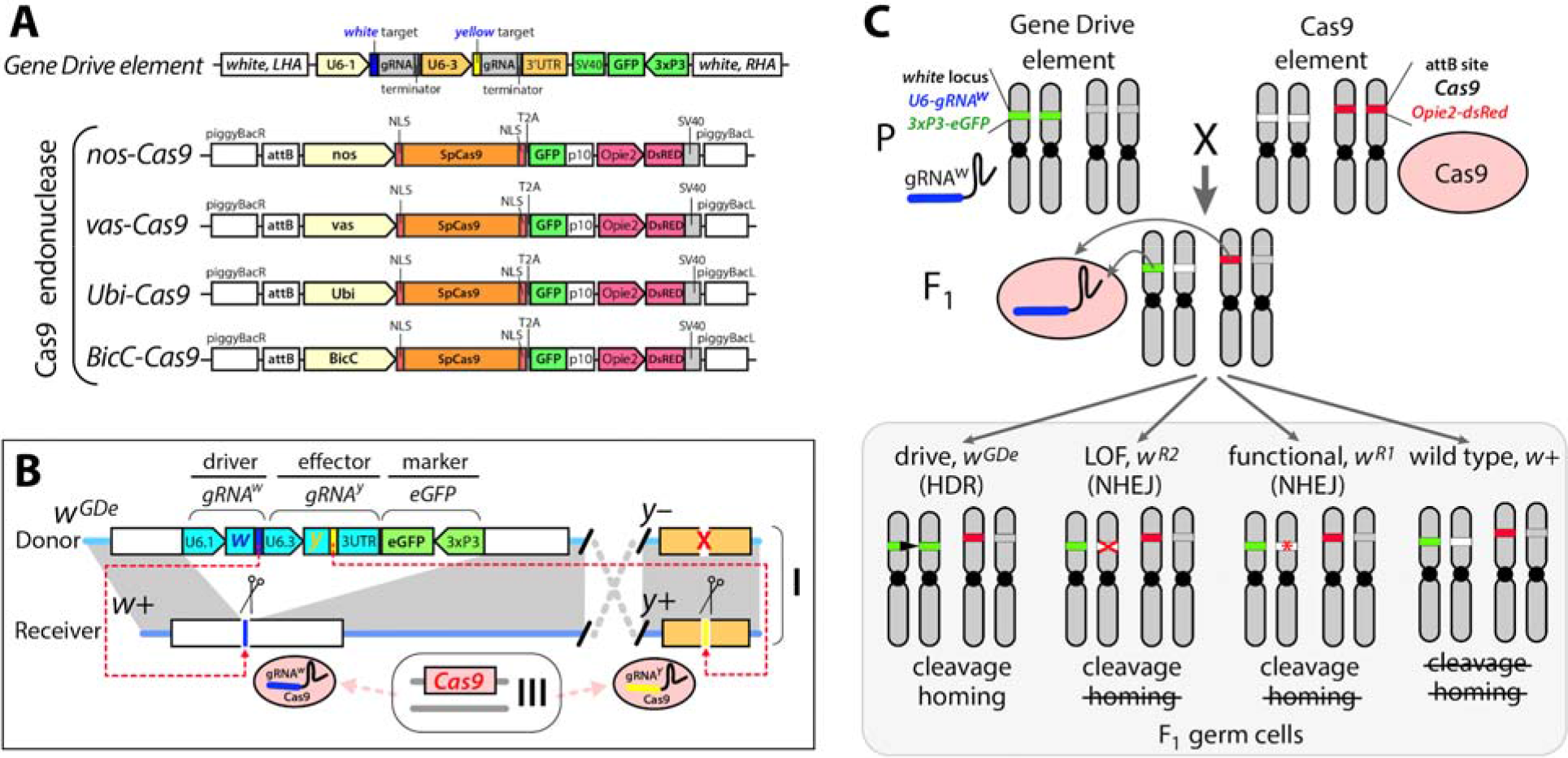
Development of the CRISPR/Cas9-mediated split-HGD system. The HGD system was split into two components: *Gene Drive* element (*GDe*) and *Cas9* endonuclease (Cas9). (**A**) Schematic maps (not to scale) of genetic constructs used to assemble split-HGD systems. The GDe contains two guide RNAs (gRNAs) targeting the DNA cleavage at *white* and *yellow* loci, and an eye-specific marker (*3xP3-GFP)* all surrounded by Left and Right Homology Arms (LHA and RHA) complementary to the *white* cut site. Four Cas9 constructs expressing *SpCas9* (Cas9) in early germline cells with *nanos* (*nos*) and *vasa* (*vas*) promoters, in late germ cells with *Bicaudal C* (*BicC*) promoter, and in both germ and somatic cells with *Ubiquitin 63E* (Ubi) promoter were integrated at the same appB site on the 3^rd^ chromosome. To track Cas9 expression, its coding sequence was linked to *eGFP* via a self-cleaving *T2A* sequence. Cas9 constructs also carry a body specific marker of transgenesis (*Opie2-DsRed*). (**B**) GDe was site-specifically inserted at *white* locus on X chromosome in *Drosophila* via HDR-mediated integration, *w*^*GDe*^. In the presence of Cas9, *GDe* direct cleavage at both *white* and *yellow* loci and can home at *white* locus from the carrier allele into a naive allele via HDR in heterozygotes (Fig. 1A). (**C**) Each element is inactive on its own and can be maintained as a homozygous parental line (P). The cross between the homozygous lines results in 100% trans-heterozygous F_1_ progeny that carry both elements. gRNA^w^ expressed by *GDe* directs cleavage at *white* locus by Cas9, which can be repaired in three different ways: via HDR using GDe as a repair template and result in homing of *GDe*; and via Non-Homologous End Joining (NHEJ) and lead to *white* loss-of-function resistance (*w*^*R2*^) allele or an in-frame functional resistance (*w*^*R1*^) allele (Fig. S4).

**Supplementary Fig. S2.**
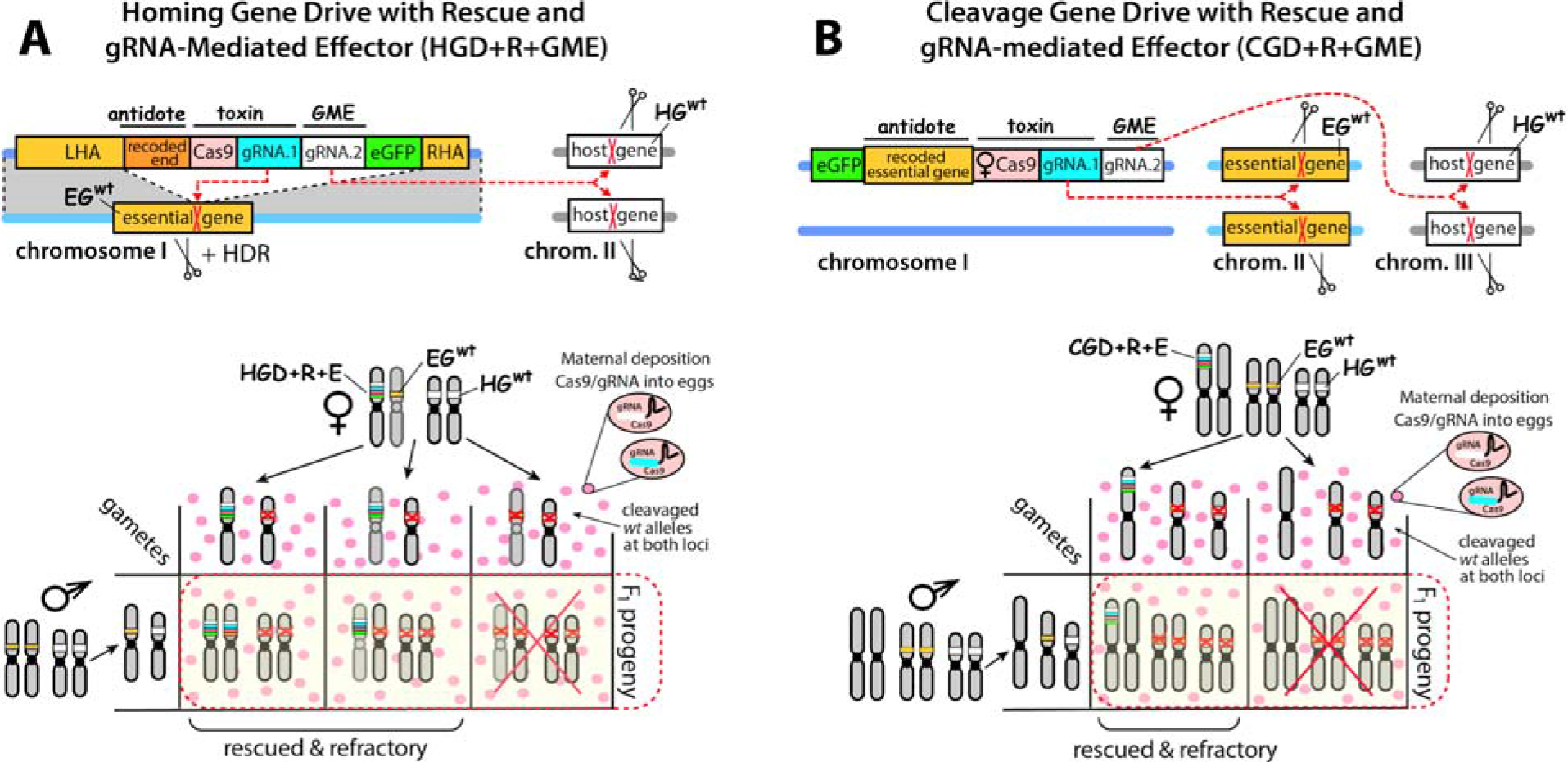
gRNA-mediated Effector (GME) incorporated into two novel gene drive designs mechanistically based on lethal biallelic mosaicism. (**A**) Schematic of Homing Gene Drive targeting an essential gene with a recoded Rescue and GME (HGD+R+GME). The HGD+R+GME expresses Cas9 and two gRNAs targeting an essential gene (EG) and host gene (HG), a marker gene (eGFP), and the cleavage-resistant recorded portion of the essential gene that is being targeted by the gRNA/Cas9 complex (rescue), which can rescue the knockout phenotype, flanked by Left and Right Homology Arms (LHA and RHA). Mechanistically, once HGD+R+GME is integrated precisely inside the EG it will direct cleavage of the EG^wt^ allele on a receiver chromosome, and induce knockout mutations that will either result in lethal biallelic mosaicism, or convert the receiver chromosome into EG^HGD+R+GME^ via homology directed repair (HDR). This ensures that only the progeny that inherit EG^HGD+R+GME^ survive, while all progeny that inherit a cleaved allele perish due to non-rescued lethal mosaicism. In addition, the HGD+R+GME induces knockout of HG located on another (or the same) chromosome, leading to desired phenotype (i.e. pathogen resistance) to its carriers. The Punnett square below depicts the genetics of how HGD+R+GME achieves a 100% transmission rate and refractoriness in F_1_ progeny. Female heterozygous for HGD+R+GME maternally deposits Cas9/gRNA complexes into every oocyte knocking out both EG and HG, and only zygotes that inherit the HDR+R+GME would survive as F_1_ progeny. Notably, HDR will convert EG^wt^ alleles into EG^HGD+R+GME^ alleles and further increase numbers of surviving F_1_ progeny and this non-Mendelian inheritance rate will depend on homing efficiencies. (**B**) Schematic of Cleavage-only Gene Drive targeting essential gene with recoded Rescue and GME (CGD+R+GME). The CGD+R+GME expresses Cas9 with multiple gRNAs targeting an ES (gRNA.1) and HG (gRNA.2), a marker gene (eGFP), and the cleavage-resistant recorded essential gene (recue) integrated at a separate genomic location from the target gene. Mechanistically, a CGD+R+GME drive relies exclusively on cleavage with no HDR required for biased inheritance. A Punnett square depicts the genetics of how CGD+R+GME achieves 100% transmission and infection resistance rates in F_1_ progeny. The female heterozygous for CGD+R+GME deposits Cas9/gRNA complexes into every oocyte, only the half of the zygotes that inherit the CDR+R+MGE in a mendelian fashion survive as F_1_ progeny, while the other half that do not inherit CDR+R+GME perish due to lethal biallelic mosaicism.

**Supplementary Fig. S3.**
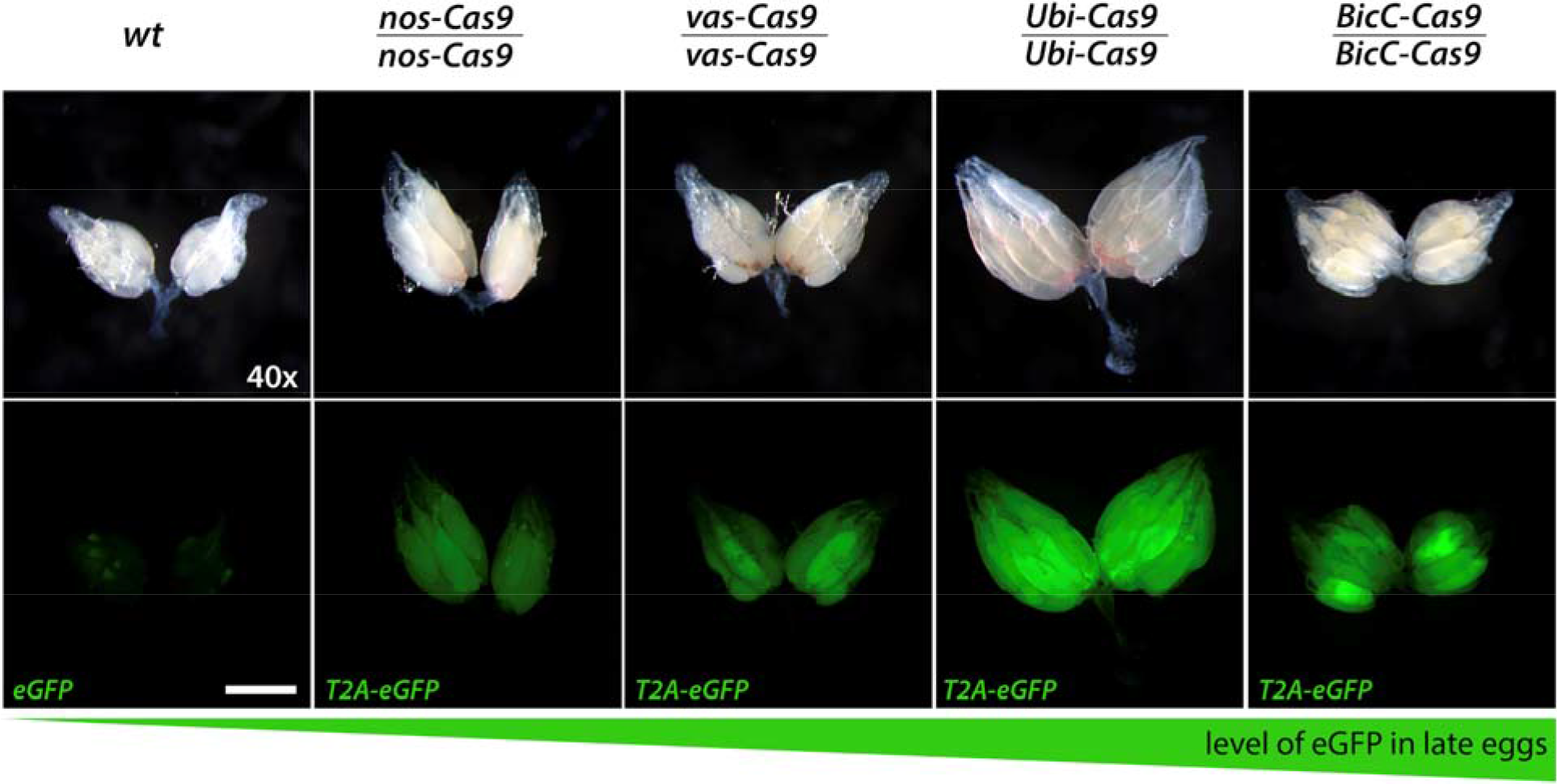
Fluorescent microscopy imaging of relative amount of maternally deposited Cas9-T2A-eGFP protein in four homozygous lines expressing *Cas9* under different promoters. Nanos (*nos-Cas9*), vasa (*vas-Cas9*), Ubiquitin-63E (*Ubi-Cas9*), and Bicaudal C (*BicC-Cas9*) constructs (Fig. S1A) were inserted at the same site on the 3rd chromosome using φC31-mediated integration. A self-cleaving T2A-eGFP sequence, which was attached to the 3’-end of Cas9 coding sequence, provided an indicator of Cas9 expression (Fig. S1A). Expression levels of eGFP in ovaries of a homozygous female from each Cas9 line were compared to that in wild type (*wt*) ovaries. Both *nos-Cas9* and *vas-Cas9* supported weak maternal deposition, while *Ubi-Cas9* and especially *BicC-Cas9* resulted in strong maternal deposition in developing eggs. Out of four tested Cas9 promoters, *nanos* and *Bicaudal C* supported the weakest and the strongest, respectively, maternal deposition into developing late eggs. Scale bars correspond to 500 μm.

**Fig. S4.**
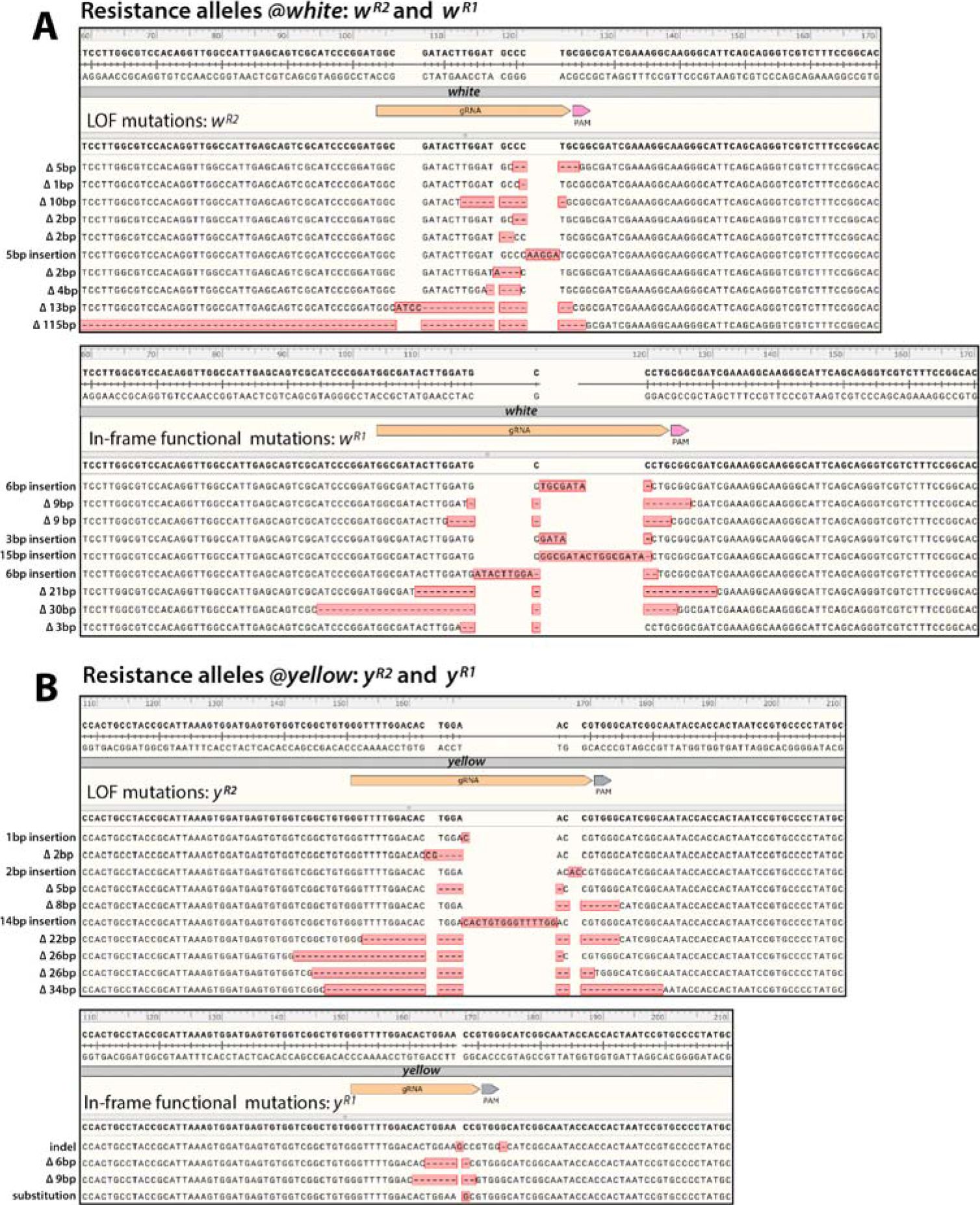
Examples of *white* and *yellow* resistance alleles generated by Cas9/gRNA-mediated DNA cleavage. Not every DSB induced by Cas9/gRNA in germ cells is repaired by HDR resulting in homing. NHEJ pathway also ligates DSBs and can lead to base insertions or deletions (*indels*) incorporated at the ligated DSBs. These *indels* change recognition sequence for gRNA and can result in mutations that are resistant to further cleavages by the same Cas9/gRNA system. We identified both types of resistance alleles – loss-of-function (LOF) (R2) and in-frame functional (R1) mutations – generated at both *white* and *yellow* loci. Diversities of *w*^*R2*^ and *w*^*R1*^ (**A**), and *y*^*R2*^ and *y*^*R1*^ (**B**) found at *white* and *yellow* loci, respectively. Homozygous LOF mutations of both *white* and *yellow* genes are viable and fertile in *Drosophila*, and thus frequencies of resistance alleles increased between F_2_ and F_3_ generations (Fig. 1D).

